# Competition for glutamate between NMDA and AMPA receptors prevents runaway synaptic dynamics

**DOI:** 10.1101/2020.04.10.035337

**Authors:** Qingchen Guo

## Abstract

Homeostatic plasticity is an important guarantee for proper neural function. However, long term potentiation (LTP) was thought of as positive feedback in Hebbian plasticity. In this condition, synapse after potentiation is prone to be further potentiated. This can cause runaway dynamics of synapse and affect the stability of neural network. In order to prevent runaway synaptic dynamics, negative feedback is needed. Upon induction of LTP, the α-amino-3-hydroxy-5-methyl-4-isoxazolepropionic acid (AMPA) receptors was increased in the postsynaptic density (PSD). Due to the competition for glutamate between AMPA receptors and N-Methyl-D-aspartic acid (NMDA) receptors, the number of opened NMDA receptor channels will reduce. Since the induction of LTP is NMDA receptors dependent, reduction of the number of activated NMDA receptors will increase the threshold of LTP induction. So the LTP of synapse itself can form a negative feedback to LTP induction. To test this hypothesis, a synaptic model with NMDA receptors and AMPA receptors coexisted was developed. When the number of AMPA receptors was increased in the PSD, the number of opened NMDA receptors was reduced though the same number of glutamate was released from presynaptic terminal. This will increase the threshold of further LTP induction and stability of synapse and neural network.

## Introduction

Long term change of synaptic transmission has a crucial role in learning and memory (1). A widely accepted model of learning and memory is Hebbian plasticity: LTP and LTD (2). LTP and LTD induction are NMDA receptors and activity dependent (3). NMDA receptors are calcium permeable(4). Either LTD or LTP can be induced by the same pattern of stimulation depending on the level of depolarization of the postsynaptic neuron (5). The level of direction of long term plasticity (6, 7). Correlated activity of presynaptic terminal and postsynaptic neuron is needed for induction of long term plasticity (8, 9). A synapse undergo potentiation may have higher probability to lead to postsynaptic spikes and increase the correlated activity of presynaptic terminal and postsynaptic neuron. This synapse is prone to be further potentiated. This form a kind of positive feedback. The long term plasticity may induce runaway synaptic dynamics (10–12). But in real nervous system, the neuron and network activity are kept relatively stable (13, 14).

Homeostatic plasticity is mechanisms to keep the synaptic and neuronal network dynamics within a computationally optimal range(13). Both local and overall homeostatic regulation have been observed (15). A lot of mechanisms have been discovered for the homeostatic plasticity(11). But the mechanism of why the synaptic weight won’t get out of control with Hebbian plasticity is still not clearly described.

To study the property of synapse after LTP induction, a model of single synapse was developed. The results indicate LTP itself can form a negative feedback to the threshold of LTP induction. This result is conflicted with previous thought that LTP is positive feedback and can cause runaway synaptic dynamics. When LTP was induced in a synapse, the number of AMPA receptors in the PSD was increased. Since AMPA receptors and NMDA receptors can both bind glutamate, they will compete for transmitter. Then the number of opened NMDA receptor channels will decrease. The threshold of LTP induction will be increased. This can prevents runaway synaptic dynamics.

## Materials and Methods

A model of glutamatergic synapse with AMPA receptors and NMDA receptors coexisted in the PSD was developed. The model consist of the glutamate release from presynaptic terminal, the diffusion of glutamate in the synaptic cleft and the interaction of glutamate and postsynaptic receptors. The model was run in the GNU Octave software. The parameters used in the model are list in Table1 (16, 17).

**Table 1.** Parameters used in the model

| Parameter | Symbol | Value |
| --- | --- | --- |
| Glutamate diffusion coeff. | D | 0.25$\mu\text{m}^2/\text{ms}$ |
| Uptake affinity(terminal) | | 2$\mu\text{M}$ |
| Uptake capacity(terminal) | $V_{\text{max}}$ | 160$\mu\text{M} / \text{ms}$ |
| Uptake rate(glia) |  | 60/ms |
| AMPA receptors density | | 800/$\mu\text{m}^2$ |
| NMDA receptors density | | 1200/$\mu\text{m}^2$ |
| AMPA receptors binding | $K_{\text{on1}}$ | 0.00918/$\mu\text{M}/\text{ms}$ |
| | $K_{\text{off1}}$ | 8.52/ms |
| | $K_{\text{on2}}$ | 0.0568/$\mu\text{M}/\text{ms}$ |
| | $K_{\text{off2}}$ | 6.52/ms |
| | $K_{\text{on3}}$ | 0.00254/$\mu\text{M}/\text{ms}$ |
| | $K_{\text{off3}}$ | 0.0914/ms |
| | $\alpha$ | 1.8/ms |
| | $\beta$ | 8.5/ms |
| | $\alpha_1$ | 0.0784/ms |
| | $\beta_1$ | 5.78/ms |
| | $\alpha_2$ | 0.01454/ms |
| | $\beta_2$ | 0.344/ms |
| | $\alpha_3$ | 0.008/ms |
| | $\beta_3$ | 0.0354.ms |
| | $\alpha_4$ | 0.3808/ms |
| | $\beta_4$ | 0.0336/ms |
| NMDA receptors binding | $k_{\text{on}}$ | 0.00918/$\mu\text{M}/\text{ms}$ |
| | $k_{\text{off}}$ | 0.0067/ms |
| | $\alpha$ | 0.20/ms |
| | $\beta$ | 0.06/ms |
| | $k_{\text{d1}}$ | 0.01/ms |
| | $k_{\text{d2}}$ | 0.01/ms |
| | $k_{\text{r1}}$ | 0.002/ms |
| | $k_{\text{r2}}$ | 0.002/ms |

### Glutamate release and diffusion

The release of transmitter from presynaptic terminal is due to the fusion of synaptic vesicle with the presynaptic membrane (18). This fusion can form a fusion pore (19). The transmitter in the synaptic vesicle can diffuse out from this fusion pore. For simplicity, the fusion pore is simulated as a point source and located at the center of the presynaptic membrane. The decay kinetics of glutamate in fused vesicle is taken as an exponential function. The time constant of glutamate concentration decrease τ in the vesicle is set to 100μs.

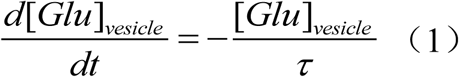

The synaptic cleft was simulated by a flat cylinder with a height of 20nm and radius of 300nm (*z, r*). The diffusion coefficient of glutamate in the synaptic cleft is 0.25μm^2^/ms (17).The diffusion function of glutamate is reduced to two dimensional due to the symmetry of cylinder.

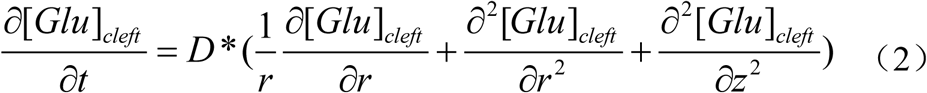

The synaptic cleft are surrounded by glia cell, so the glutamate cannot diffuse out. The glia cell and the presynaptic terminal can uptake glutamate (20, 21).

### The kinetics of glutamate and receptor interaction

When glutamate diffused to the postsynaptic region, glutamate can bind to postsynaptic receptors. The binding kinetics of glutamate to NMDA receptors and AMPA receptors are adopted from Holmes’s model(17). The kinetics are schemed in Fig. 1. Both the AMPA receptors and NMDA receptors have two binding sites of glutamate. The receptors can be opened or desensitized by the binding of glutamate. The density of AMPA receptors and NMDA receptors are 800/μm^2^ and 1200/μm^2^ separately (16). Both AMPA receptors and NMDA receptors are evenly distributed in the PSD. For simplicity, the block of NMDA receptors by magnesium is not considered.

**Fig. 1.**
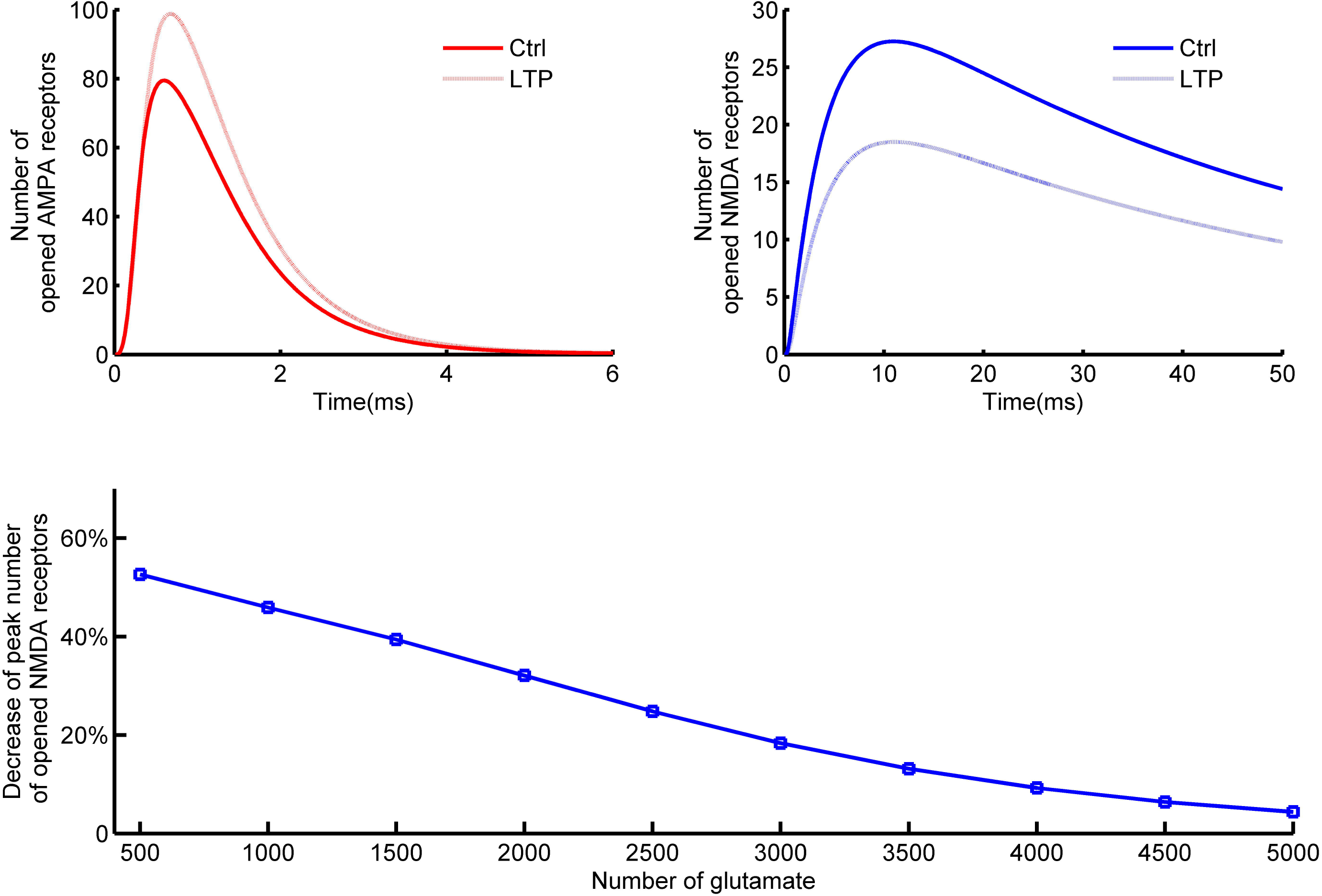
The kinetics of glutamate and NMDA receptors and AMPA receptors interaction. (*A*) The kinetics of glutamate binding to NMDA receptors. NMDAR represents NMDA receptor, Glu represents glutamate. NMDAR * indicate the opened state of NMDA receptor channel and NMDAR_D is desensitized state of NMDA receptor. The block of NMDA receptors by magnesium is not considered. (*B*) Kinetics of glutamate binding to AMPA receptors. AMPAR represents AMPA receptors, AMPAR* is the opened state of AMPA receptor channel and AMPAR_D is the desensitized state of AMPA receptor.

## Results

### Non-saturation of receptors and competition between NMDA receptors and AMPA receptors for glutamate

Both AMPA receptors and NMDA receptors are glutamate receptors (22). The number of transmitter is variable per vesicle and the postsynaptic response of AMPA receptors can range from several picoamps to more than one hundred picoamps (23, 24). So the glutamate receptors can sense variable concentration of glutamate. It is useful to determine whether glutamate can saturate the postsynaptic receptors and the relation between the response of receptors and number of glutamate released from presynaptic terminal. Using the synaptic model, the relationship between the number of glutamate released from presynaptic terminal and synaptic response was tested. The response of AMPA receptors and NMDA receptors increase following the increase of glutamate released from presynaptic terminal. But the increment of the response of receptors get smaller with the same amount of increment of glutamate due to the saturation of receptors (Fig.2*A*). The estimated number of glutamate per vesicle is about 2000 (25). The postsynaptic receptors are not saturated by this concentration of glutamate. This indicate that single package of glutamate cannot saturate postsynaptic receptors. This result is consist with previous reports that single package of glutamate cannot saturate postsynaptic receptors (26, 27). The glutamate sensitivity and affinity are different between AMPA receptors and NMDA receptors (Fig.2*B*).

**Fig.2.**
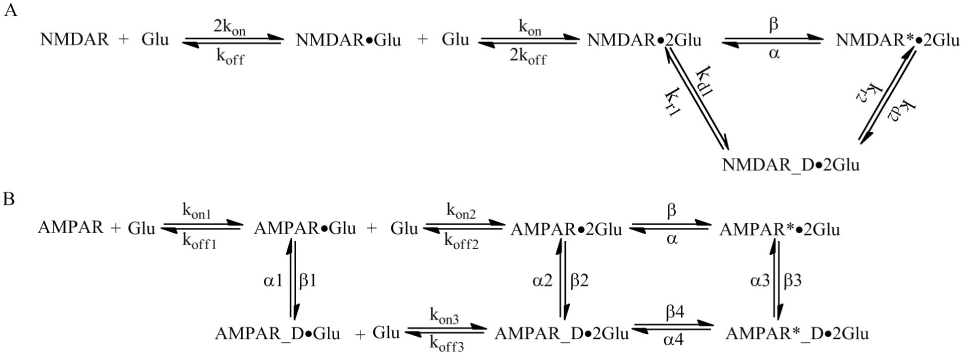
Non-saturation of receptors and competition between NMDA receptors and AMPA receptors for glutamate. (*A*) The relation of the number of opened AMPA receptors and NMDA receptors and the number of glutamate released from presynaptic terminal with these two types of receptors coexist. The red line is the peak number of opened AMPA receptor channels, the blue line is the peak number of opened NMDA receptor channels. (*B*) Normalized relation of opened AMPA receptors and NMDA receptors and the number of glutamate released from presynaptic terminal with these two types of receptors coexist. (*C*) Glutamate sensitivity of AMPA receptors without the affection of NMDA receptors. The solid line is the peak number of opened AMPA receptor channels with AMPA receptors and NMDA receptors coexist. The dotted line is the peak number of opened AMPA receptor channels with only AMPA receptors in the PSD. (*D*) Glutamate sensitivity of NMDA receptors without the affection of AMPA receptors. The solid line is the peak number of opened NMDA receptor channels with AMPA receptors and NMDA receptors coexist. The dotted line is the peak number of opened NMDA receptor channels with only NMDA receptors in the PSD.

In the PSD, AMPA receptors and NMDA receptors coexist and they are both glutamate receptors. Since single package of glutamate cannot saturate postsynaptic receptors, these two types of receptors may compete for glutamate. The response of one type of receptors can be affected by the other. To study how these two types of receptors affect each other, only one type of receptors was placed in the postsynaptic side in the model. The relation between the number of glutamate released from presynaptic terminal and the number of opened receptor channels with only one type of receptors placed in the PSD was obtained. Compared to the condition of two types of receptors coexist, the response of just single type of receptors is bigger (Fig.2*C, D*). Since the density of AMPA receptors is larger than NMDA receptors, the response of NMDA receptors are more severely affected by AMPA receptors.

### Negative feedback of Hebbian plasticity

LTP and LTD are cellular processes that are involved in learning and memory. AMPA receptors and NMDA receptors coexist in the PSD and both of them can undergo long term plasticity (28, 29). But long term plasticity of AMPA receptors is mostly widely observed. Only the long term plasticity of AMPA receptors was considered in this work. Since AMPA receptors and NMDA receptors are both glutamate receptors. Increasing one type of receptors can reduce the response of the other. How long term plasticity of AMPA receptors will affect the response of NMDA receptors? In the model, the density of AMPA receptors was increased one fold to simulate the effect of LTP. The response of the AMPA receptors was increased almost one fold (Fig.3*A*). But the response of NMDA receptors was decreased due to the competition for glutamate between AMPA receptors and NMDA receptors (Fig.3*B*). So LTP can enhance the synaptic weight, but the response of NMDA receptors was reduced. The activation of NMDA receptors is necessary for induction of long term plasticity. This indicate induction of LTP can increase the threshold of further LTP induction. So long term plasticity can form negative feedback to itself. This will prevent runaway synaptic dynamics. This mechanism may play an important role in homeostatic plasticity of synapse. To test how the density of AMPA receptors change affect the response of NMDA receptors, the density of NMDA receptors was kept at 800/μm^2^ and the density of AMPA receptors ranged from 500/μm^2^ to 5000/μm^2^. Increasing the density of AMPA receptors, the response of AMPA receptors increased nearly linearly. While the response of NMDA receptors was decreased following the increase of the density of AMPA receptors (Fig.3*C*).

**Fig.3.**
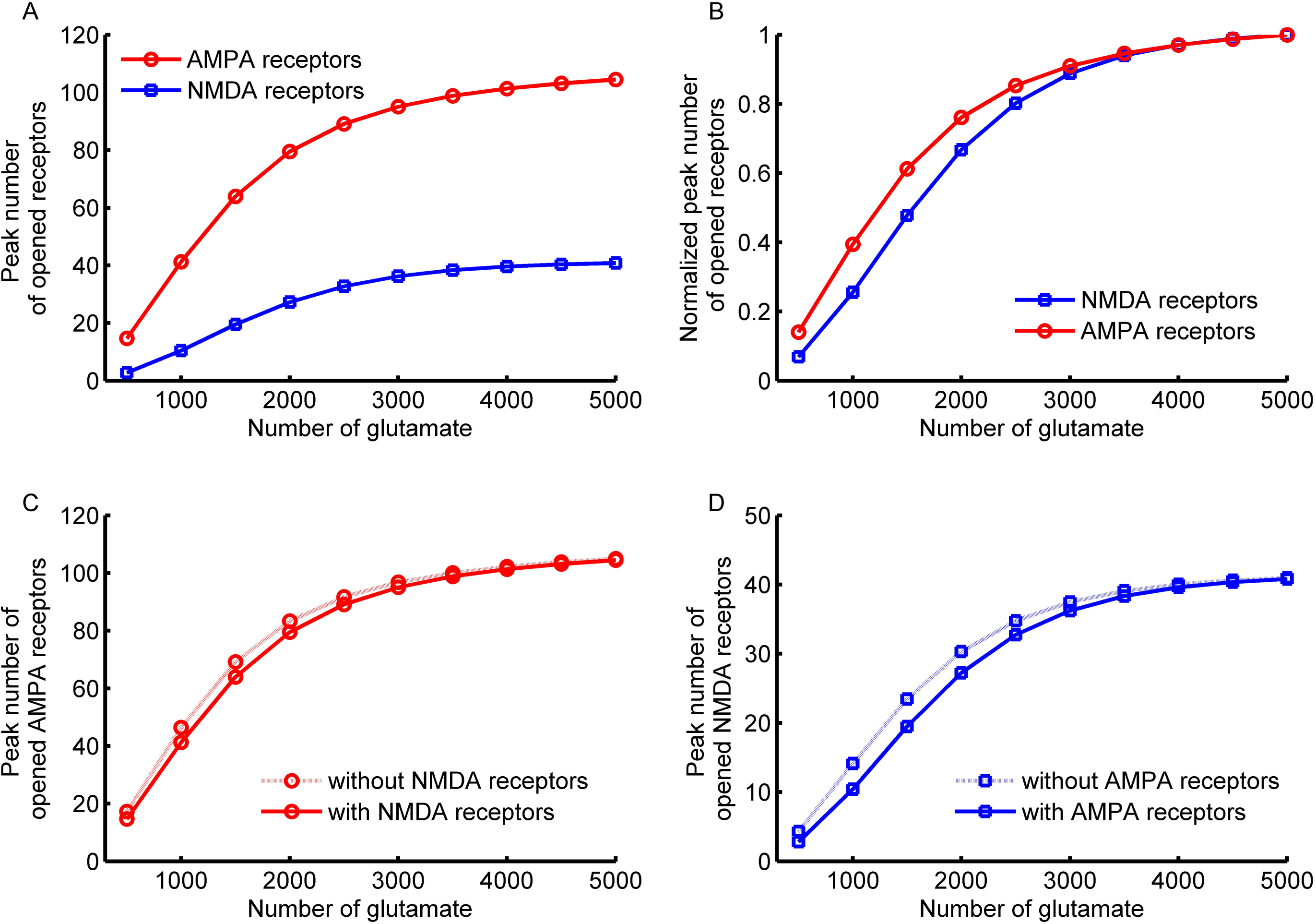
Negative feedback of LTP induction. (*A*) The response of AMPA receptors before and after LTP of AMPA receptors. The solid line is the response of AMPA receptors before LTP and the dotted line is the response of AMPA receptors after LTP (increase the density of AMPA receptors one fold). (*B*) The response of NMDA receptors before and after LTP of AMPA receptors. The solid line is the response of NMDA receptors before LTP of AMPA receptors, the dotted line is the response of NMDA receptors after LTP of AMPA receptors. (*C*) Relation between peak number of activated AMPA receptors and NMDA receptors and the density of AMPA receptors when the density of NMDA receptors was kept unchanged. The red line is the peak number of opened AMPA receptors, the blue line is the peak number of opened NMDA receptors.

The negative feedback of LTP is due to the competition of AMPA receptors and NMDA receptors for the same transmitter. So glutamate concentration released from presynaptic terminal will affect the weight of this negative feedback. Physiologically, the number of single package of transmitter are variable and can be regulated. How the quantal size will affect the negative feedback effect of long term plasticity? To test this effect, the glutamate number per vesicle ranged from 500 to 5000 per vesicle in the model. Increasing the number of glutamate reduced the difference of the response of NMDA receptors between the control one and the LTP of AMPA receptors (Fig.4*B*). When the glutamate can totally saturate postsynaptic receptors, the competition between AMPA receptors and NMDA receptors for glutamate get negligible.

**Fig.4.**
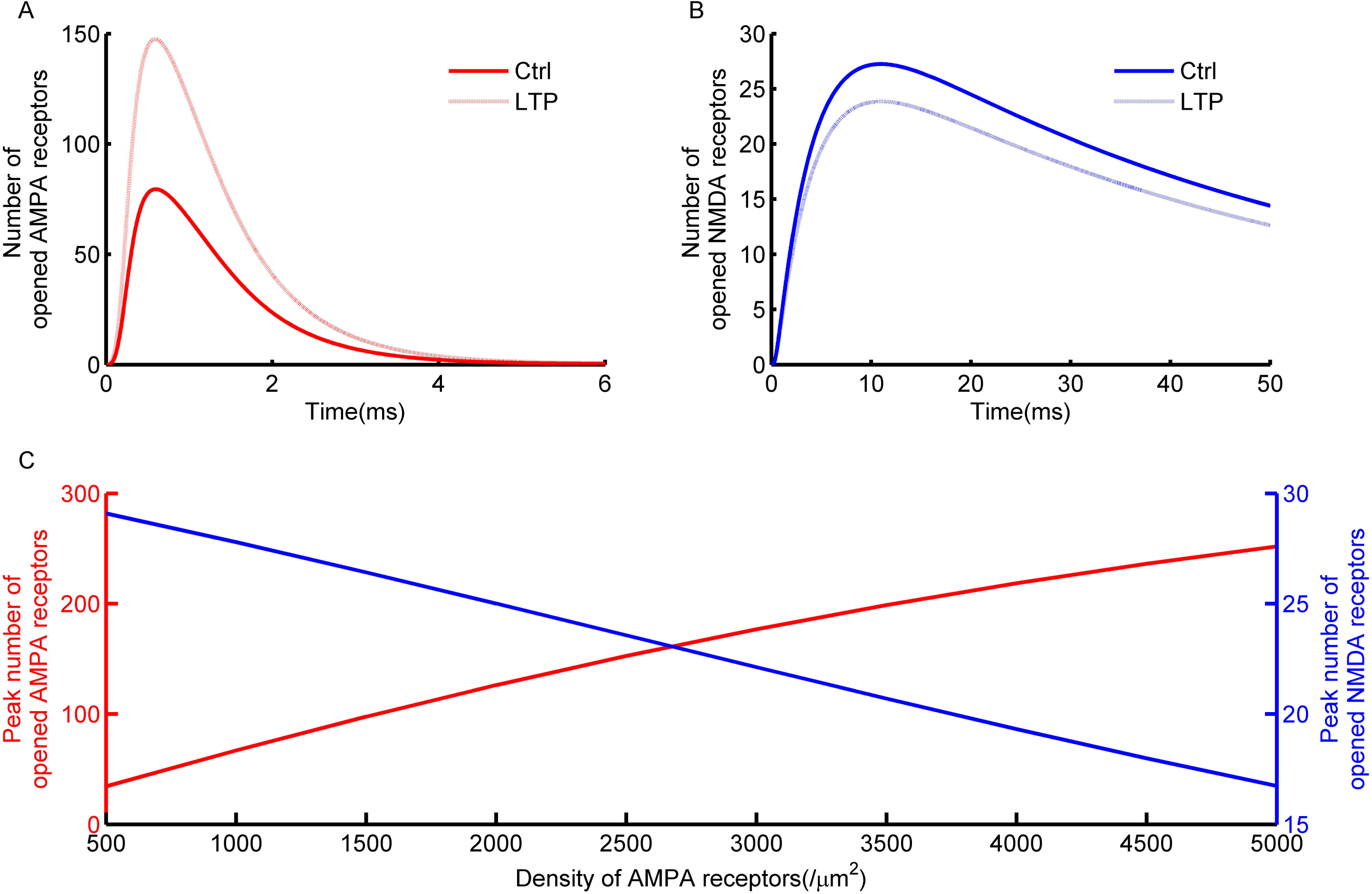
The weight of negative feedback of LTP is affected by glutamate concentration. (*A*) Sample traces of the response of NMDA receptors before and after LTP of AMPA receptors with different numbers of glutamate released from presynaptic terminal. (*A1*) 1000 glutamate per vesicle, (*A2*) 2000 glutamate per vesicle, (*A3*) 3000 glutamate per vesicle. (*B*) Dependence of the reduction of the response of NMDA receptors on the number of glutamate released from presynaptic terminal.

There are evidences that the NMDA receptors and AMPA receptors are non-homogeneously distributed in the PSD (30–32). AMPA receptors are denser at the periphery of the PSD and the density of NMDA receptors is highest at the center of the PSD. To test how the different distributions of AMPA receptors and NMDA receptors affect the competition for glutamate. The distribution of NMDA receptors are set as a gaussian distribution with the highest density at the center of PSD and the standard deviation is 0.1μm. The distribution of AMPA receptors is also gaussian distribution with the highest density located at 0.15μm to the center of PSD and the standard deviation is half of the distribution of NMDA receptors (Fig.5*A*). The results show even the distribution of AMPA receptors and NMDA receptors are different, the competition for glutamate are still exist. After the LTP of AMPA receptors, the response of NMDA receptors was reduced (Fig.5*C*).

**Fig.5.**
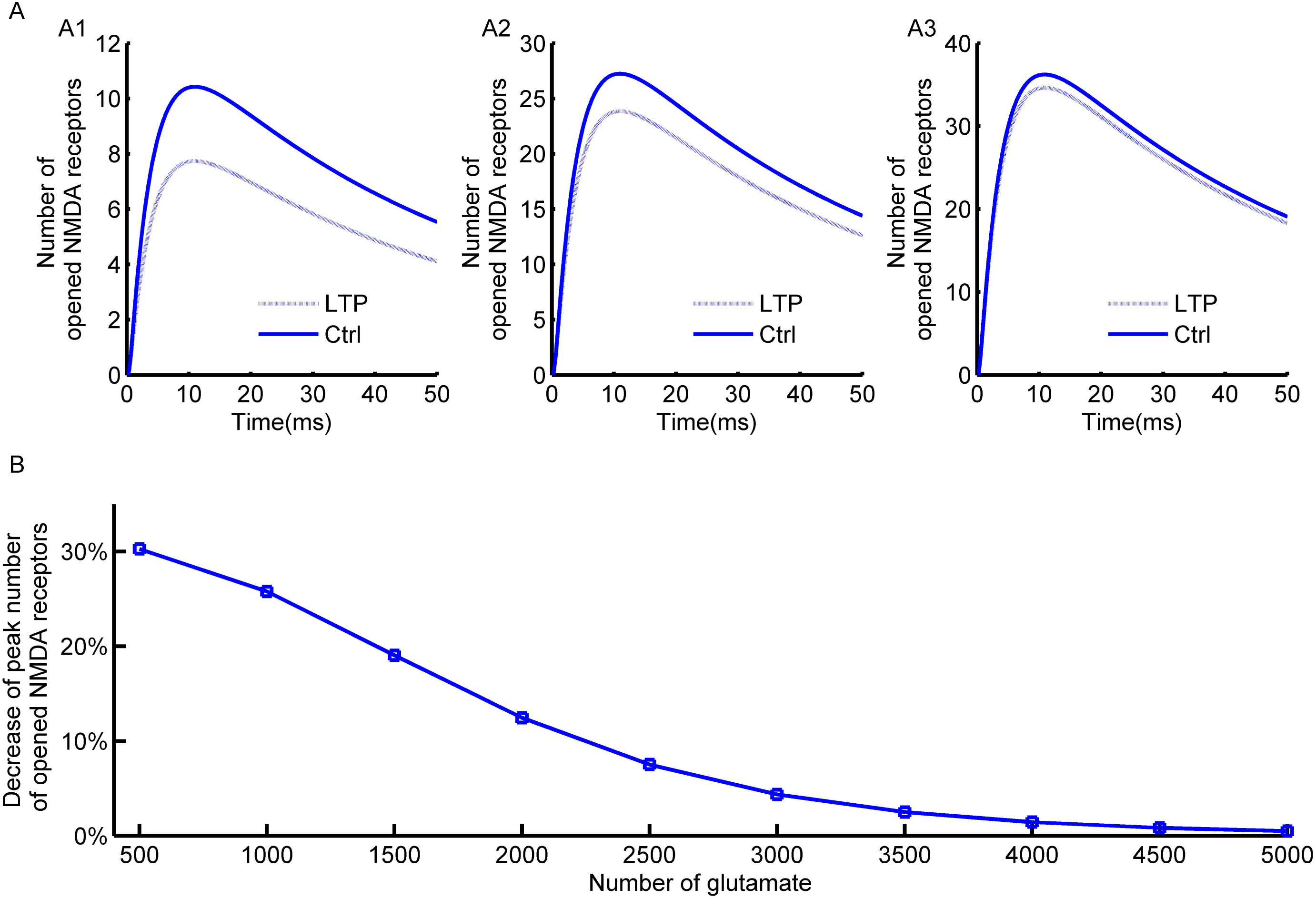
Non-homogenous distribution of AMPA receptors and NMDA receptors does not eliminate the competition for glutamate. (*A*) The distribution of AMPA receptors and NMDA receptors in the PSD. (*B*) The response of AMPA receptors before and after LTP induction (increase the density of AMPA receptors one fold). (*C*) The response of NMDA receptors before and after the LTP of AMPA receptors.

### Synaptic enlargement enhance negative feedback of LTP

Following synaptic strength enhancement after LTP induction, the structure of synapse will also change (33, 34). To simulate this effect, the area of the PSD was increased one fold in the model. The density of AMPA receptors and the total number of NMDA receptors remained unchanged. Since the area of the PSD was increased, the density of NMDA receptors was decreased as well. The result show the decrement of the response of NMDA receptors after LTP of AMPA receptors was larger compared to the density change of AMPA receptors (Fig.6*A*). But the increment of the response of AMPA receptors after LTP of AMPA receptors is smaller than in the condition of increasing the density of AMPA receptors in the PSD (Fig.6*B*). One reason is synaptic cleft was also enlarged, so the concentration of glutamate was reduced in the synaptic cleft. Another is the density of NMDA receptors was reduced due to the enlargement of PSD. So the enlargement of synapse will enhance the negative feedback of long term plasticity of AMPA receptors. The decrement of the response of NMDA receptors is also glutamate concentration dependent in this condition (Fig.6*C*). Increasing the number of glutamate released from presynaptic terminal decreased the difference of the response of NMDA receptors between the control one and LTP of AMPA receptors. When glutamate can totally saturate postsynaptic receptors, the competition between AMPA receptors and NMDA receptors get negligible.

**Fig.6.**
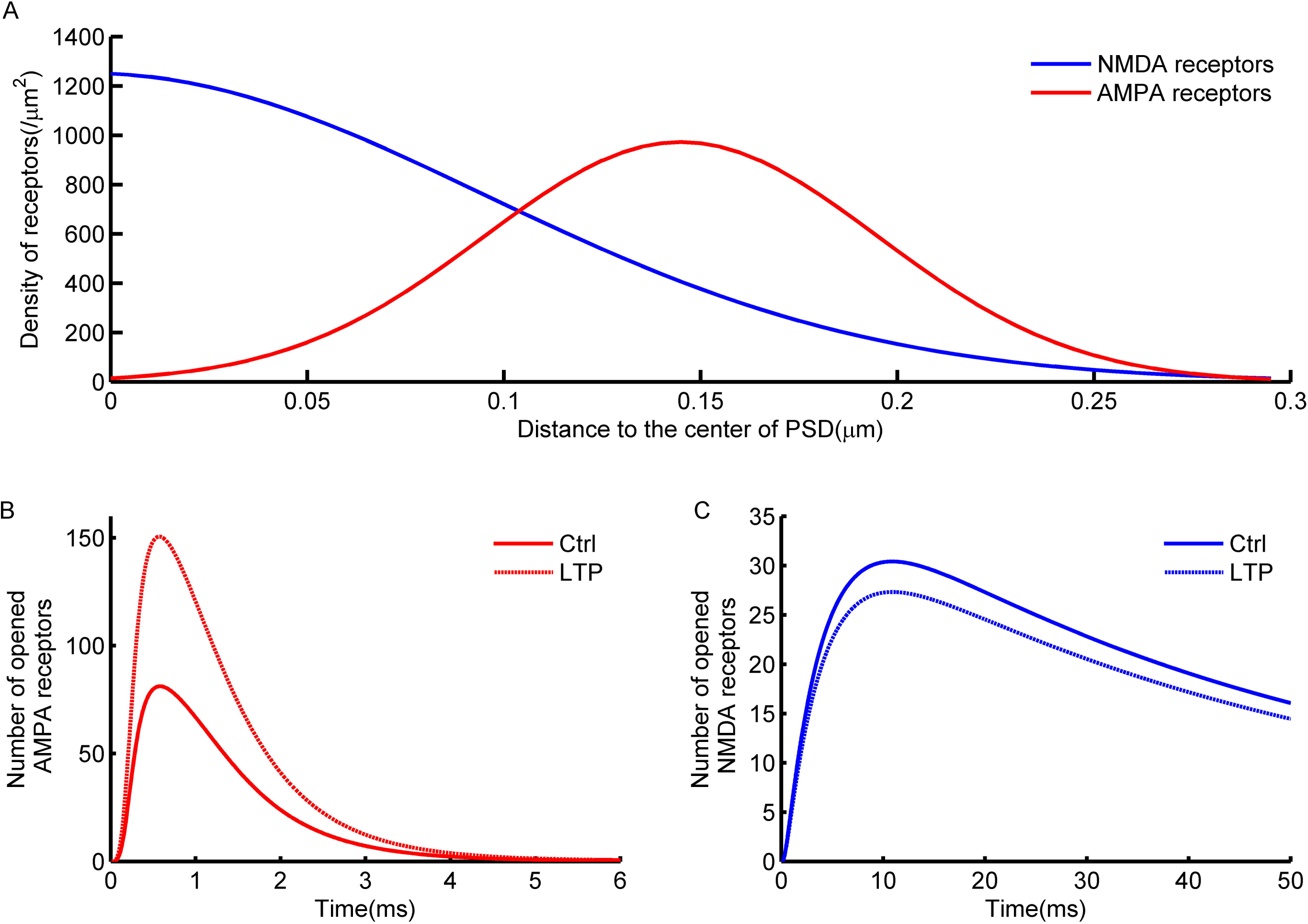
The effect of synaptic structure change after LTP induction. (*A*) The response of AMPA receptors before and after LTP of AMPA receptors. The solid line is the response of AMPA receptors before LTP and the dotted line is the response of AMPA receptors after LTP (the area of PSD was increased one fold, the density of AMPA receptors and the total number of NMDA receptors was not changed). (*B*) The response of NMDA receptors before and after LTP of AMPA receptors. The solid line is the response of NMDA receptors before LTP of AMPA receptors, the dotted line is the response of NMDA receptors after LTP of AMPA receptors. (*C*) The dependence of the reduction of the response of NMDA receptors on the number of glutamate released from presynaptic terminal.

## Discussion

A great diversity of plasticity has been discovered in nervous system and they are likely governed by independent mechanisms (35). Homeostatic plasticity is an important mechanism to preserve stable electrophysiological properties of neuron. In recent years, important progress has been made toward identifying molecules and signaling processes required for homeostatic forms of neuroplasticity. A new mechanism of homeostatic plasticity was provided in this work. In glutamatergic synapse, the AMPA receptors and NMDA receptors can affect each other due to the competition for the same transmitter. It is usually thought that Hebbian plasticity is positive feedback and can cause runaway synaptic dynamics. This work indicate that LTP is a negative feedback to the threshold of LTP induction. Induction of LTP can increase the threshold of further LTP induction. This negative feedback is due to the competition for glutamate between AMPA receptors and NMDA receptors. This may be a new mechanism of homeostatic plasticity and can prevents runaway synaptic dynamics.

The competition of AMPA receptors and NMDA receptors for glutamate is based on the result that the postsynaptic receptors are not saturated by glutamate released from presynaptic terminal. If the glutamate can saturate postsynaptic receptors, the competition effect of NMDA receptors and AMPA receptors will be eliminated. By now there are a lot of experimental results show the postsynaptic receptors are not saturated by transmitter (27, 36, 37). Functionally, no saturation of receptors can provide more dynamic space for the regulation of synaptic function. So it is reasonable to think that the postsynaptic receptors are not saturated by moderate activity of synapse.

An important factor that affect the negative feedback of LTP is the number of glutamate released from presynaptic terminal. In the model, the number of glutamate per vesicle is adopted from Rusakov’s estimate (25). The real number of glutamate per vesicle is still not determined. But it is confirmed that the number of glutamate are variable and can be regulated by the activity of synapse (38, 39). If the number of glutamate released from presynaptic terminal is smaller, the negative feedback of LTP will get larger.

The negative feedback of LTP to the threshold of LTP induction is based on that only the number of AMPA receptors in the PSD was increased after induction of LTP. There is report that the induction of long term potentiation will increase NMDA receptors and AMPA receptors proportionally. But the increase of NMDA receptors is slower than AMPA receptors (40). The proportional change of AMPA receptors and NMDA receptors can eliminate the effect of negative feedback of LTP. But there are also evidences conflicted with the proportional increase of AMPA receptors and NMDA receptors in the LTP induction of AMPA receptors. In nucleus accumbens, induction of calcium dependent LTP of non-NMDA receptors will produce a simultaneous LTD of NMDA receptors (41). This will enhance the negative feedback of LTP. In the dentate gyrus, the NMDA receptors increase during development, but the ratio of NMDA receptors to AMPA receptors decrease (42). In hippocampus during development, the silent synapse acquired AMPA receptors with little change of NMDA receptors (43). There is also evidence that the NMDA receptors mediated current decrease while the response of AMPA receptors increase at an auditory synapse during development (44). So it is reasonable to think that NMDA receptors receptor will not increase after LTP induction of AMPA receptors. LTP of AMPA receptors can reduce the response of NMDA receptors, also LTD of AMPA receptors can increase the response of NMDA receptors. So the overall ratio of the response of AMPA receptors and NMDA receptors may still keep constant.

